# ACE: adaptive cluster expansion for maximum entropy graphical model inference

**DOI:** 10.1101/044677

**Authors:** J. P. Barton, E. De Leonardis, A. Coucke, S. Cocco

## Abstract

**Motivation:** Graphical models are often employed to interpret patterns of correlations observed in data through a network of interactions between the variables. Recently, Ising/Potts models, also known as Markov random fields, have been productively applied to diverse problems in biology, including the prediction of structural contacts from protein sequence data and the description of neural activity patterns. However, inference of such models is a challenging computational problem that cannot be solved exactly. Here we describe the adaptive cluster expansion (ACE) method to quickly and accurately infer Ising or Potts models based on correlation data. ACE avoids overfitting by constructing a sparse network of interactions sufficient to reproduce the observed correlation data within the statistical error expected due to finite sampling. When convergence of the ACE algorithm is slow, we combine it with a Boltzmann Machine Learning algorithm (BML). We illustrate this method on a variety of biological and artificial data sets and compare it to state-of-the-art approximate methods such as Gaussian and pseudo-likelihood inference.

**Results:** We show that ACE accurately reproduces the true parameters of the underlying model when they are known, and yields accurate statistical descriptions of both biological and artificial data. Models inferred by ACE have substantially better statistical performance compared to those obtained from faster Gaussian and pseudo-likelihood methods, which only precisely recover the structure of the interaction network.

**Availability:** The ACE source code, user manual, and tutorials with example data are freely available on GitHub at https://github.com/johnbarton/ACE.

**Supplementary information:** Supplementary data are available

## 1 Introduction

Interpreting patterns of correlations in data is a fundamental problem across scientific disciplines. A common approach to this problem is to infer a simple graphical model that explains the statistics of the data through a network of effective interactions between the variables, which may then be used to generate new predictions [1]. The goal of this approach is to disentangle the direct interactions between variables from their correlations, which arise through a combination of direct and indirect effects. Here we focus on a particular family of undirected graphical models, referred to as Potts models in the language of statistical physics, which have recently been applied to study a wide variety of biological systems. Applications include inference of the effective connectivity of populations of neurons, and their patterns of firing activity, based on data from multi-electrode recordings [2, 3, 4, 5], and the prediction of protein contact residues [6] and the fitness effects of mutations [7, 8, 9] based on the analysis of multiple sequence alignments (MSAs).

Unfortunately, the inference of Potts models from data is challenging. The computational time required for naive Potts inference algorithms scales exponentially with the system size, rendering the problem intractable for realistic systems of interest. Various approximations have been employed to combat this problem, including Gaussian and mean-field inference [10], perturbative expansions [11, 12], and pseudo-likelihood methods [13, 14]. These approximate methods can successfully capture the general structure of the network of interactions, recovering, in particular, contact residues in the three-dimensional structure of protein families [6, 15, 16, 17, 18, 19, 20], but the resulting models typically give a less accurate statistical description of the data [21]. Alternately, algorithms based on iterative rounds of Monte Carlo simulation [22, 8, 23] are capable of inferring models that accurately reproduce the observed correlations, but they are typically slow to converge.

Here we describe an extension of the adaptive cluster expansion (ACE) method, originally devised for binary (Ising) variables [24, 25], to more general (Potts) variables taking multiple categorical values. We also describe new computational methods for faster inference, including a fast Monte Carlo learning procedure and the optional incorporation of prior knowledge about the structure of the interaction graph. The algorithm has been successfully applied to real data with as many as several hundred variables, including studies of neural activity in the retina and prefrontal cortex [24, 25, 5, 26], human immunodeficiency virus (HIV) fitness based on protein MSA data [8, 27], and lattice protein models [28]. Below we illustrate the application of this method to both real and artificial data sets. We show that models inferred by ACE give an excellent reconstruction of the statistics of the data. They also accurately recover, considering sampling limitations, true underlying model parameters when they are known, and can achieve comparable performance to state-of-the-art methods for predicting structural contacts in protein family data. We compare these results to those obtained using other approximate inference methods, focusing in particular on pseudo-likelihood methods.

### 1.1 Background

The Potts model emerges naturally in the statistical description of complex systems. Consider a system of *N* variables described by the configuration х = 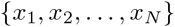, with 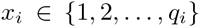. The number of discrete categories *q_i_* that each variable *x_i_* can take on, which we refer to as states, may depend on the variable index *i*. For proteins the states correspond to particular amino acids, while for neurons they represent the binary (firing or silent) state of activity. Given a set of measurements of the system, the empirical average over the sampled configurations gives us the 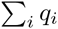 individual and 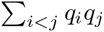 pairwise frequencies for the different states of each variable in the data. We denote the individual and pair-wise frequencies by *p_i_*(*a*) and *p_ij_*(*a, b*), respectively, where *i, j* are the index of the variables and *a, b* are the index of the states. As an example, x could represent sequences in a MSA, with *p_i_*(*a*) the frequency of the amino acid labeled by *a* in column *i* of the alignment, and *p_ij_*(*a, b*) the frequency of the pair of amino acids *a, b* in columns *i, j*.

The simplest, or maximum entropy [29], probabilistic model capable of reproducing the observed frequencies is a Potts model, which assigns a probability to every configuration of the system x:

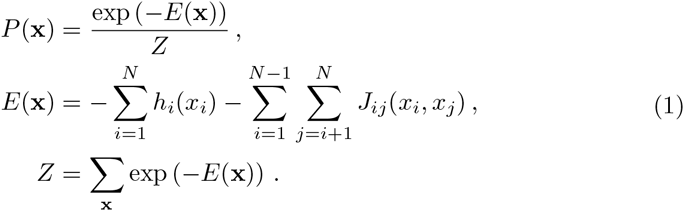

Here the partition function *Z* is a normalizing factor which ensures that all probabilities sum to one. In the simple case that all the variables *x_i_* are binary, this model is referred to as an Ising model. The parameters *h_i_*(*a*) and *J_ij_*(*a, b*) in the energy function *E*, called fields and couplings, must be chosen such that variable averages (correlations) in the model match those in the data, i.e.

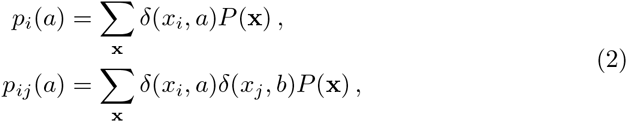

where *δ* is the Kronecker delta function. The problem of finding the parameters *h_i_*(*a*)*, J_ij_*(*a,b*) that satisfy Equation (2) is referred to as the inverse Potts problem. Note that the probability of any configuration remains unchanged under the transformation of the couplings and fields given by 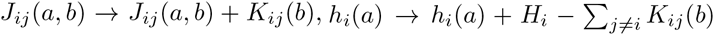 for any *K*. This “gauge invariance” reduces the number of free parameters in the Potts model to *q_i_* – 1 fields for each site and (*q_i_* – 1)(*q_j_* – 1) couplings for each pair of sites.

Formally, the inverse Potts problem is solved by the set of fields and couplings that maximize the average log-likelihood or equivalently, those that minimize the cross-entropy between the data and the model,

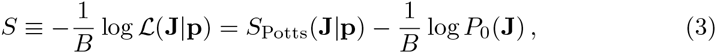

where *B* is the number of data points in the sample, and

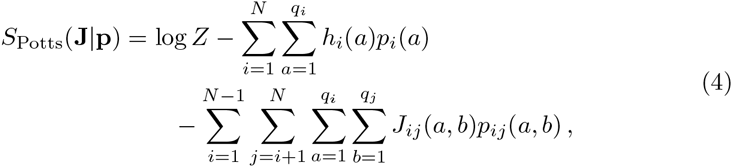

and *P*_0_ is a prior distribution for the parameters. Here for simplicity we have written the set of all individual and pairwise variable frequencies as **p** and the set of all fields and couplings as **J**. Note that, ignoring the contribution of the prior distribution, the cross-entropy S is equivalent to the entropy of the inferred model satisfying Equation (2).

The inclusion of a prior distribution helps to avoid overfitting, while also improving convergence. A Gaussian prior distribution for the parameters is a typical choice, which contributes a term

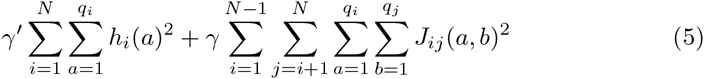

to Equation (3). The addition of this factor ensures that the solutions of the inverse problem are not at plus or minus infinity. Note that this form of the regularization is not invariant under gauge transformations. Thus, the results of the inference including the regularization do have some dependence on the gauge choice. Other forms of regularization are also possible (see Supplementary Materials). Note that the presence of the partition function *Z* in Equation (4) precludes direct numerical maximization of the likelihood when the system size is large, since this requires summing over all 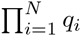 configurations of the system. Alternate methods of solving the inverse Potts problem involve approximation schemes or rely on computationally costly Monte Carlo simulations, as described above.

## 2 Methods

### 2.1 Adaptive cluster expansion

The adaptive cluster expansion [24, 25] is based on the formal decomposition of the regularized cross-entropy Equation (3) into a sum of contributions from subsets (or clusters) of the variables 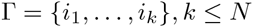,

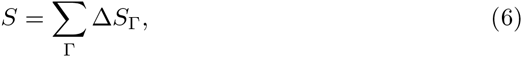

where the sum is over all nonempty subsets of the *N* variables. The terms ∆*S*_Γ_, referred to as cluster entropies, are recursively defined,

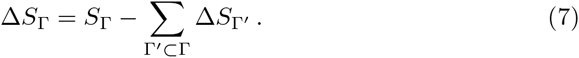

Here *S*_Γ_ denotes the maximum of Equation (3) restricted only to the variables in Γ. Thus, *S*_Γ_ depends only on the frequencies *p_i_*(*a*), *p_ij_*(*a, b*) with *i*, *j* ∊ Γ Provided that the number of variables in Γ is small (typically ≤ 20) numerical maximization of the likelihood restricted to Γ is tractable. Note that the definition of ∆*S*_Γ_ ensures that the sum over all clusters Γ in Equation (6) yields the log-likelihood for the entire system of *N* variables.

Neglecting the regularization term, the single variable cluster contributions are the entropies of the variables taken as if they were independent, 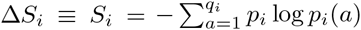. The two variable entropy is 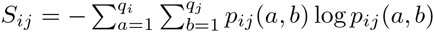 (see Supplementary Materials for more details). The cluster entropy for a pair of variables is then 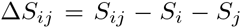, which is equivalent to the mutual information. It is zero when *p_ij_*(*a,b*) *= p_i_*(*a*)*p_j_*(*b*), i.e. when the two variables are independent. In general, ∆*S*_Γ_ is a measure of the inter-dependence between the variables in the cluster which cannot be accounted for by smaller clusters.

The main idea of this approach is to approximate the cross-entropy (and simultaneously, the parameters that maximize it) by limiting the sum in Equation (6) to a restricted set of clusters Γ that give the most important contributions to it. As shown in [24, 25], neglecting clusters with small contributions to the cross-entropy helps to avoid overfitting. We define a threshold *t* on the cross-entropy to separate the significant clusters from those which can be neglected. Starting from a large value of the threshold (typically *t* = 1), such that only a few clusters are selected, the algorithm proceeds through two nested iterations. The outer loop is on the value of the threshold *t*, which is progressively lowered until enough clusters are included to yield a model consistent with the data. The inner loop constructs the set of clusters Γ with contributions to the cross-entropy 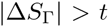 and yields an approximation of the cross-entropy and the model parameters at the threshold *t*. Contributions to the cross-entropy from clusters within the same interaction sub-network partially compensate, and thus summing up clusters according to 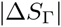 allows for a faster convergence of Equation (6) [24, 25]. The algorithm stops at the first value of the threshold *t* where the inferred model fits the sampled averages and correlations Equation (2) to within the statistical error due to finite sampling (see Section 3.2).

The algorithm for the inner loop, including the selection and summation of individual clusters, is as follows. Given a list *L_k_* of clusters of size *k*, beginning with the list of all clusters of size *k* = 2:

1. For each cluster 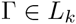
  (a) Compute *S*_Γ_ by numerical minimization of Equation (3) restricted to Γ.
  (b) Record the parameters minimizing Equation (3), called J_Γ_.
  (c) Compute ∆*S*_Γ_ using Equation (7).
2. Add all clusters 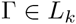 with 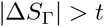 to a new list 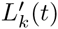.
3. Construct a list *L_k+1_* of clusters of size *k + 1* from overlapping clusters in 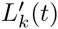.

The rule for constructing new clusters of size *k + 1* from selected clusters of size *k* can be lax (such that a new cluster Γ is added provided that any pair of size *k* subclusters, 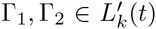 and 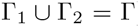) or strict (such that a new cluster is only added if all of its *k* + 1 subclusters of size *k* belong to 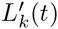). The above process is then repeated until no new clusters can be constructed.

After the summation of clusters terminates, the approximate value of the parameters minimizing the cross-entropy, given the current value of the threshold, is computed by

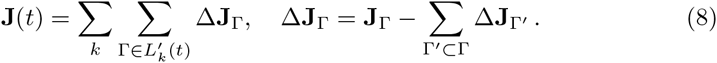

Note that this formula generally yields sparse solutions because nonzero couplings are only included in Equation (8) if some clusters containing them have been selected. In this algorithm the dominant contribution to the computational complexity often comes from the evaluation of the partition function *Z* for large cluster sizes, which requires 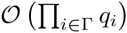 operations to compute.

### 2.2 Compression of the number of Potts states

As mentioned in Section 1.1, the number of states each variable may take on need not be the same for all variables in a system. States with zero (or otherwise very small) probabilities may be observed very infrequently in real, finitely-sampled data, and the relative error on the corresponding correlations due to finite sampling is large.

To limit overfitting and reduce the computational time, the low probability states can be effectively grouped together according to a given compression parameter. Here we present two conventions for compressed representations of the data. First, for each variable we can treat explicitly the states observed with probability larger than a cutoff value 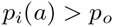 while grouping all infrequently observed values into the same state. Alternatively, we can order the states by their contribution to the total single site entropy *S_q_* and choose a reduced model in which only the first *k* states are modeled explicitly, with *k* chosen to capture a certain fraction f of the site entropy (Supplementary Materials). The final *q* – *k* states are grouped together. The frequency of the regrouped Potts state is then the sum of the frequencies of the states which have been regrouped: 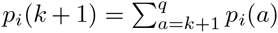. Once the reduced model is inferred, one can recover a complete model by modifying the field parameter for the regrouped states, 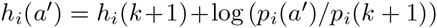, while keeping the couplings to the value of the regrouped state *J_ij_*(*a*’, *b*) *= J_ij_*(*k +* 1,*b*). For states with zero probabilities in the data, we fix the fields from the regularization alone.

### 2.3 Expansion around a reference structure

ACE is a two-fold algorithm: it builds up the interaction graph while also inferring the corresponding parameters that reproduce the correlated structure of the data. This expansion can accelerated if the interaction graph is known, or by incorporating a priori information about the interaction graph. It is also possible to expand the cross-entropy around its Gaussian approximation.

- If the list of directly interacting variables is known, one can run the expansion using this restricted set of sites such that clusters of larger size are built up only from the initial list of interacting pairs. For proteins this procedure can be applied using the real contact map, known from structural information, or alternatively the one derived with fast inference approaches such as DCA or plmDCA [6, 19]. to obtain a selected list of putative contacts and then use the cluster expansion to infer the interactions between them.
- As shown in [25] for the Ising model, one can analytically calculate the log-likelihood and the parameters that maximize it under the Gaussian approximation with an ad hoc *L_2_*-norm regularization (where the regularization strength depends on the variable frequencies). It is then possible to perform the cluster expansion around this Gaussian reference model.

### 2.4 Refinement with Boltzmann Machine Learning (BML)

In cases where convergence of the cluster algorithm alone is not sufficiently fast, it is often more expedient to use the output set of fields and couplings as starting values for a Boltzmann Machine Learning (BML) routine. In typical cases, provided that the inferred model is not too sparse, this procedure can lead to rapid convergence of the model even when the starting error is large.

Here we adapted the RPROP algorithm for neural network learning [30] to the case of Potts models. Given an input set of fields and couplings, we first compute the model correlations 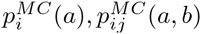 through Monte Carlo simulation. The couplings and fields are then updated according to the gradient of the log-likelihood, multiplied by a parameter-specific weight factor

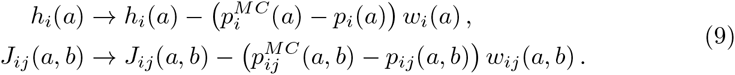

Regularization can also be incorporated by adding 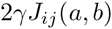, or the analogous term for fields, to the gradient. Here the weights 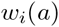 and 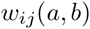 are also updated with each iteration of the algorithm. At each iteration, if the sign of 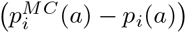 is the same as in the previous round, 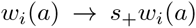, else 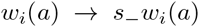, and similarly for the 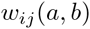. This acceleration of weight parameters allows appropriate step sizes to be chosen adaptively for each coupling and field. To prevent steps sizes from becoming too large or too small, the weight parameters are restricted to lie between some *w*_min_ and *w*_max_. Typical choices of the weight bounds and update multipliers are 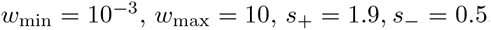. Note that we choose 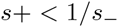 so that, if the sign of one of the terms of the gradient continually switches, the corresponding weight decreases.

## 3 Results

### 3.1 Description of test data and their preprocessing

#### 3.1.1 Potts models on Erdős-Rényi random graphs (ER05)

We consider an example of a Potts model with *q* = 21 states, where the network of interactions is described by an Erdős-Rényi random graph with *N* = 50 variables. Each edge in the interaction graph is included with probability 0.05. Field and coupling values for interacting pairs of sites are selected from a Gaussian distribution (Supplementary Materials). We compute the correlations through Monte Carlo sampling of *B* = 10^4^ configurations. In the results shown below we compressed rarely-observed Potts states with 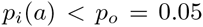 and used 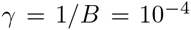, performing the inference in the gauge of the compressed Potts state.

#### 3.1.2 Lattice protein model (LP *S_B_*)

We consider an alignment of 5 x 10^4^ protein sequences with *N* = 27 sites, arranged in a 3 x 3 x 3 cube, selected according to their exactly computable [31] folding probability *S_B_* (see [28], Supplementary Materials). In the results below we have remove never-observed amino acids (i.e. compression with *p_o_* = 0), and used the regularization 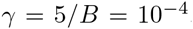. Couplings and fields corresponding to the least frequently observed amino acid at each site are gauged to zero.

#### 3.1.3 Trypsin inhibitor protein family (PF00014)

We study an alignment of 4915 sequences downloaded from the PFAM database for the trypsin inhibitor protein family (PF00014). After removing columns with > 50% gaps the number of sites is *N* = 53. We reweight the contribution of each sequence to the correlations according to its similarity to other sequences in the alignment, an approach commonly used to attenuate phylogenetic correlations [6]. Here we show results in the consensus gauge after compressing rarely-observed amino acids with 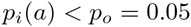, using 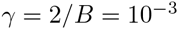. Additionally, we note that gaps in the MSA are not generally modeled well in the Potts model representation with pairwise interactions, as they tend to be present in long stretches, especially at the beginning and the end of the alignment [20]. Such stretches of highly correlated gaps slow down the inference procedure because they give rise to large clusters. Here we have processed the data to replace gaps by random amino acids with the same frequency as observed in the non-gapped sequences.

#### 3.1.4 HIV p7 nucleocapsid protein

The HIV nucleocapsid protein p7 plays an essential role in multiple aspects of viral replication [32]. We downloaded a MSA of 4131 p7 sequences from individuals infected by clade B viruses from the Los Alamos National Laboratory HIV sequence database (www.hiv.lanl.gov). After removing columns with > 95% gaps, the remaining number of sites is *N* = 71. Here we do not reweight sequences by similarity, given that they are all phylogenetically related. We have replaced gaps in the alignment as described above, compressed rarely-observed amino acids with *f_S_* = 90%, and chosen 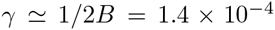. Inference is performed in the consensus gauge.

#### 3.1.5 Multi-electrode recordings of cortical neurons

We divided a 20 minute recording of the firing activity of 32 cortical neurons into a set of *B* = 1.5 x 10^5^ time bins of 10ms, treating each time window as an observation of the system. During each time window, the variable for each neuron *i* was assigned *x_i_* = 1 if the neuron was active at least once during that time, and zero otherwise. Here we take 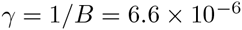.

### 3.2 Convergence of the cluster expansion algorithm

As mentioned in Section 2.1 for each threshold *t* used to select clusters in the ACE expansion, the model individual 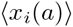 and pairwise 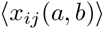 frequencies are compared to the data’s frequencies *p_i_*(*a*) and *p_ij_*(*a,b*). We define a relative error as the ratio between the deviations of the predicted observables from the data 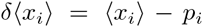 and 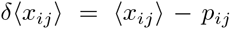 and the expected statistical fluctuations due to finite sampling: 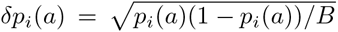, 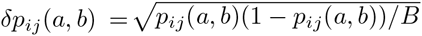. We define the normalized maximum error as

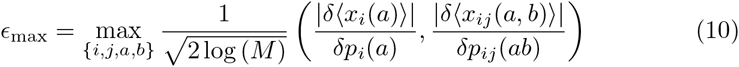

where *M* is the total number of one- and two-point correlations.

Figure 1 shows the behavior of *∊*_max_ and the cross-entropy as a function of the threshold the five data sets described above. The cross-entropy *S* approaches a constant value as the threshold is decreased. In all cases except for the lattice protein model, the algorithm converges at *∊*_max_ ~ 1, when the correlations are reproduced to within the expected error due to finite sampling. The expansion slows dramatically for the lattice protein model at a fairly high value of the threshold due to the large number of states included at each site in the model (typically *q* = 19). The computational cost of calculating the partition function is a limiting factor as the maximum cluster size increases, corresponding to *K*_max_ = 7 at the stopping point in Fig. 1. At this point, BML is needed to refine the parameters inferred through the cluster expansion. Note that, even in cases when the error appears large, convergence of the BML procedure is often rapid because only small changes to the parameters may be necessary to obtain a model that accurately reproduces the correlations.

**Figure 1:**
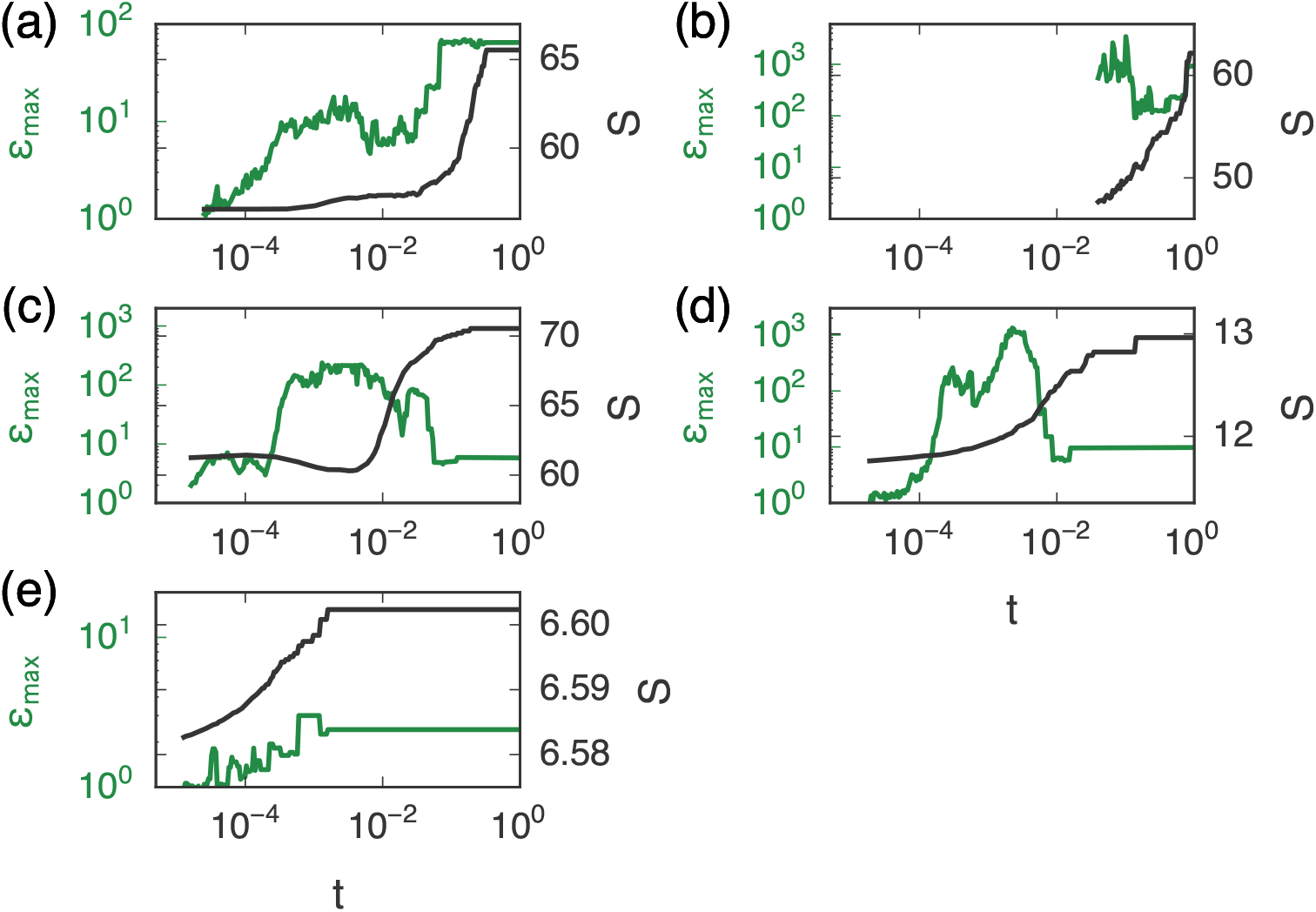
Convergence of the duster expansion as a function of the threshold *t* for (a) ER005, (b) LP *S_B_* (c) PF00014, (d) HIV p7, and (e) cortical data. As the threshold is lowered, the cross-entropy *S* approaches a constant value. In all cases except for LP *S_B_* the normalized maximum error *∊*_max_ reaches 1 through the duster expansion alone. For LP *S_B_* a Monte Carlo learning procedure is used to refine the inferred parameters and reach *∊*_max_ ≃ 1.

Convergence of the algorithm can also be more difficult for alignments of long proteins or those with very strong interactions. In such cases one may observe large oscillations in the cross-entropy as a function of the threshold, and large (≥ 10 sites) clusters may appear even at high thresholds. Strong regularization (γ > *1/B*) can help to dampen these oscillations, after which it can be returned to ≈ *1/B* during the BML procedure.

### 3.3 Parameters of the ER05 model are recovered by ACE

In Fig. 5 we show that the 2 x 10^4^ underlying parameters for the ER05 model corresponding to the explicitly modeled Potts states are accurately recovered by ACE. These states are better sampled and therefore they have smaller statistical uncertainties. In the model inferred by plmDCA, which includes no reduction in the number of states, there are around 10^6^ parameters. Those corresponding to the explicitly modeled states are recovered fairly well (with some errors in the fields), but parameters corresponding to compressed states are difficult to infer due to insufficient sampling (see Supplementary Materials for details and analysis of errors in inferred parameters due to finite sampling).

**Figure 2:**
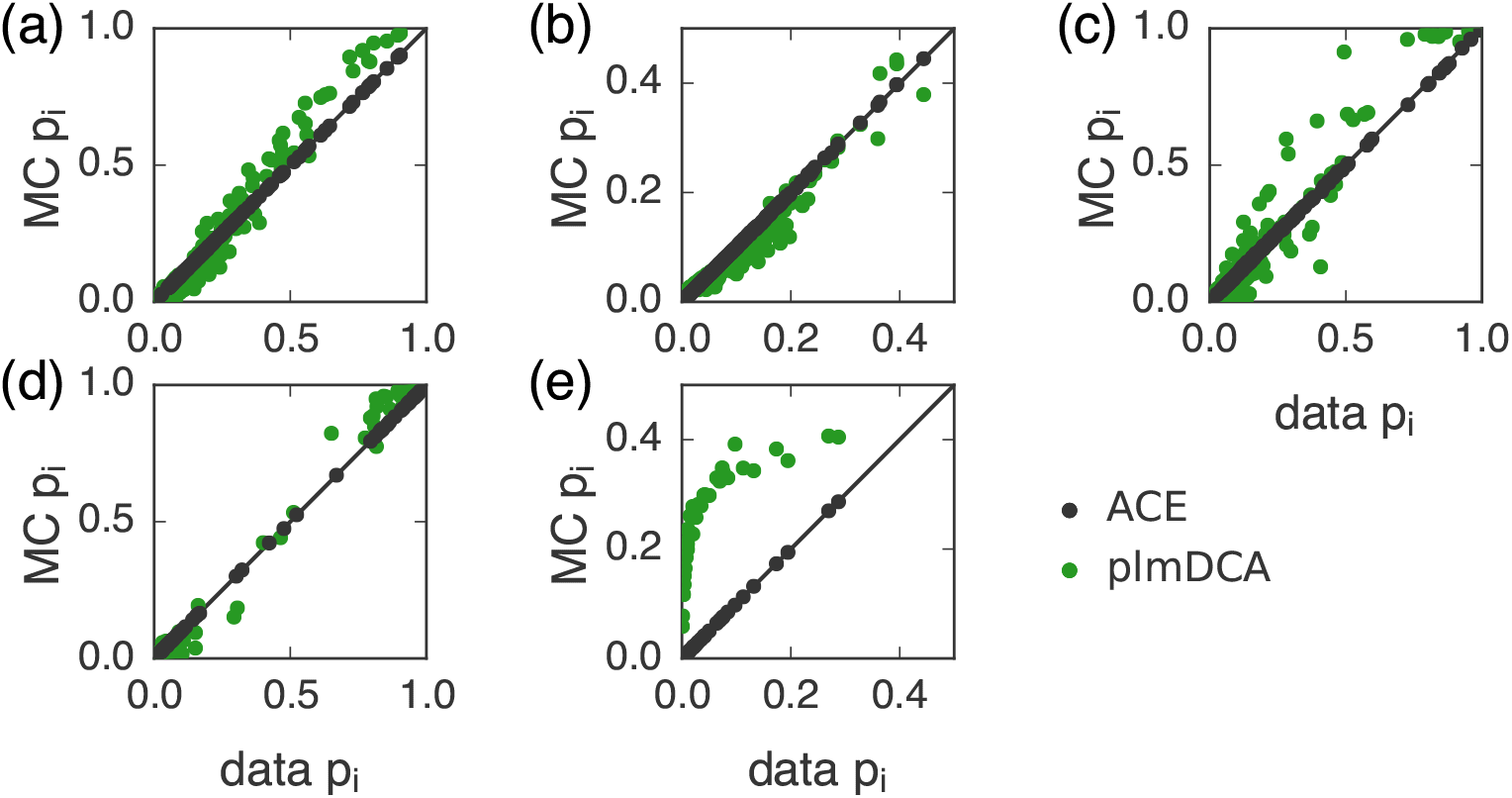
ACE outperforms plmDCA in recovering the single variable frequencies for models describing (a) ER005, (b) LP *S_B_*, (c) PF00014, (d) HIV p7, and (e) cortical activity. The results for plmDCA are obtained with the regularization γ = 0.01, which gives better results for the correlations than lower values of the regularization strength (see Supplementary Materials).

**Figure 3:**
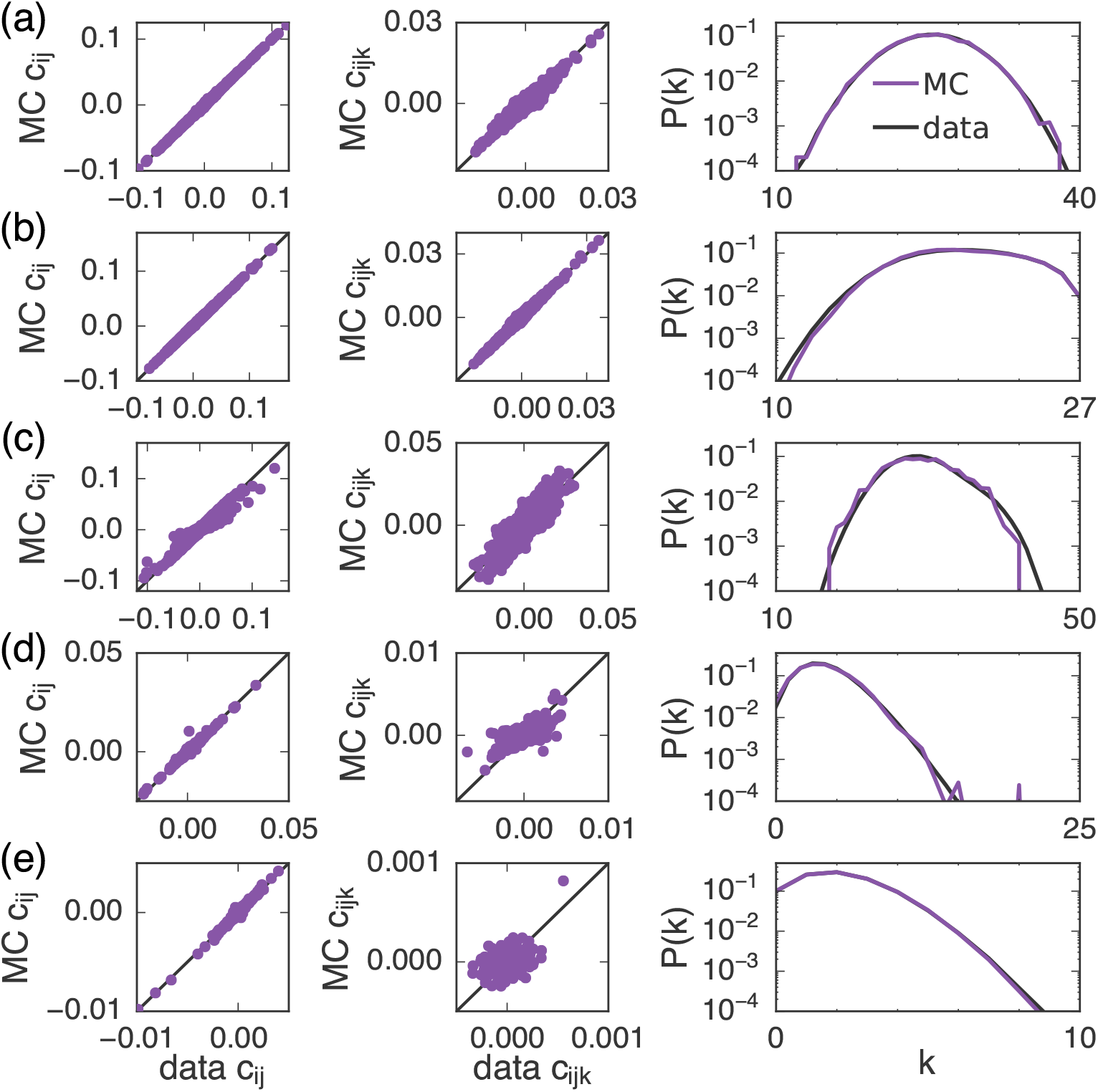
Fit for models describing (a) ER005, (b) LP *S_B_*, (c) PF00014, (d) HIV p7, and (e) cortical activity. ACE recovers the connected pair correlations *c_ij_*(*a, b*) *= p_ij_*(*a, b*) *– p_i_*(*a*)*p_j_*(*b*) (left). The inferred model also successfully captures higher order correlations present in the data, such as the connected three-body correlations (center) and the probability *P*(*k*) of observing a configuration with *k* differences from the consensus configuration (right).

**Figure 4:**
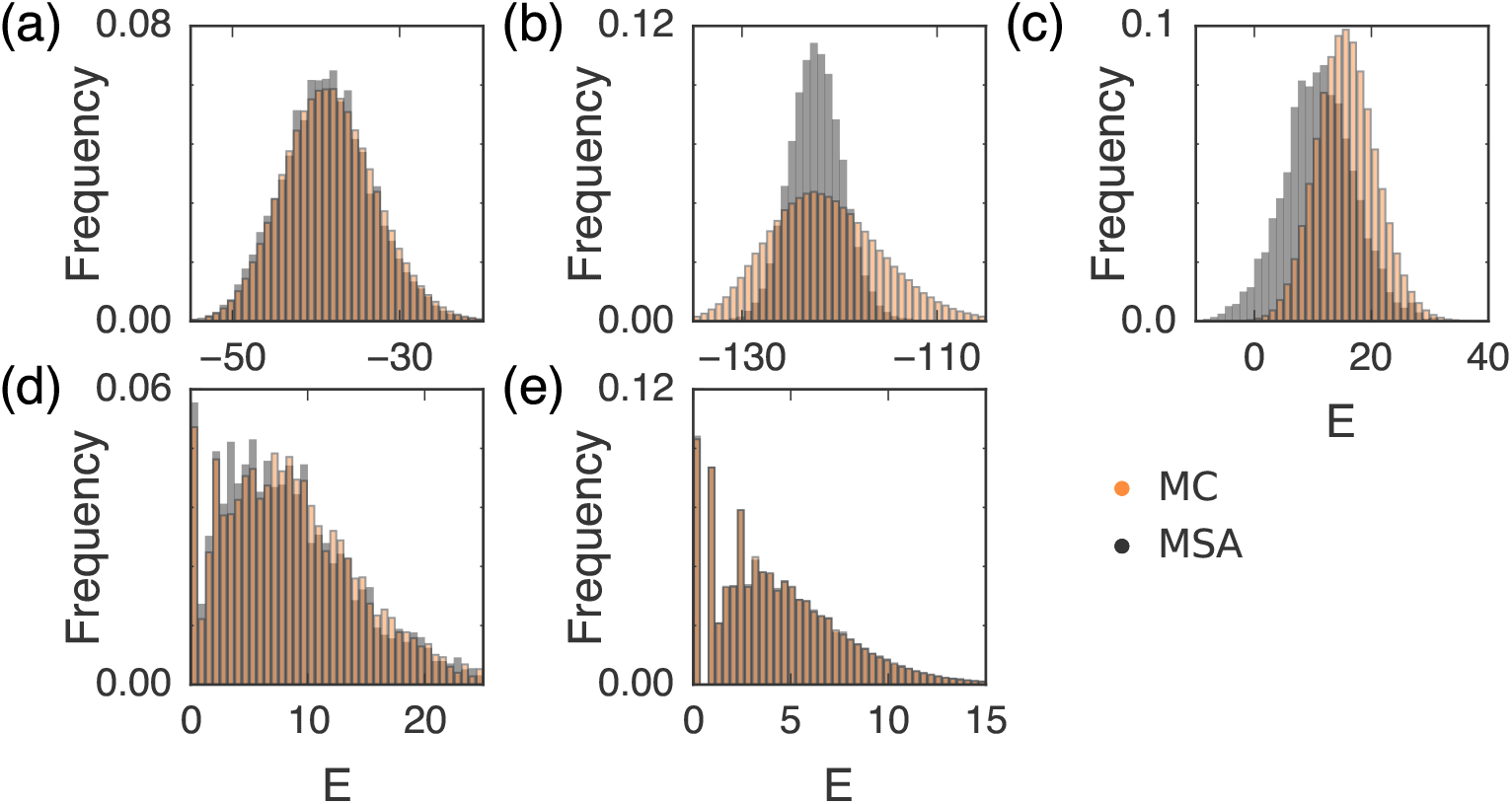
Histograms of the data (MSA) and model (MC) energy distributions for (a) ER005, (b) LP *S_B_*, (c) PF00014, (d) HIV p7, and (e) cortical activity. Monte Carlo sampling of the inferred Potts model describing each set of data yields a distribution of energies similar to the empirical distribution, a further check on the consistency of the model fit beyond the fitting of correlations.

**Figure 5:**
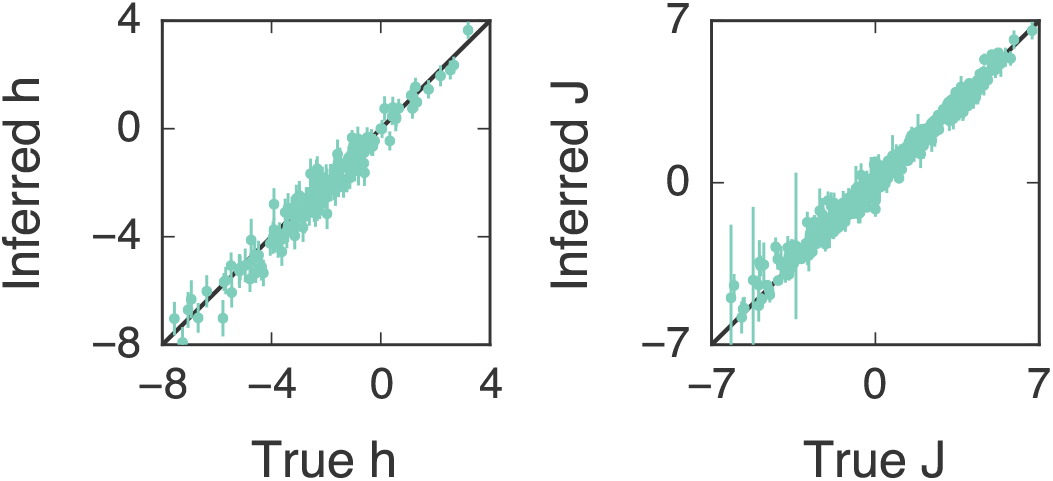
ACE accurately recovers the the true fields *h* (left) and couplings J (right) corresponding to Potts states with *p_i_*(*a*) *≥* 0.05 for the ER05 model. Error bars denote standard deviation in estimated parameters due to finite sampling.

### 3.4 Statistics of the data are accurately reproduced

Figures 2 and 3 show how the model inferred by ACE reproduces the statistics of the input data. In all cases the model accurately captures the input probabilities and pairwise connected correlations within the expected error due to finite sampling, as expected.

We also find that higher order correlations in the data can be accurately reproduced. Figure 3 shows the 3-point connected correlations and the distribution *P*(*k*) of Hamming distances *k* between the sampled configurations and the configuration in which each site takes on the most probable value (i.e. the consensus sequence for proteins). In the neural case the most probable configuration is the silent one and therefore *P*(*k*) is the probability to have *k* active neurons in the same time window. Models inferred by ACE outperforms those from plmDCA [19], see Fig. 2 and Supplementary Materials for higher order statistics.

Comparing the distribution of energies *E* for configurations sampled from the inferred model to the distribution obtained from the original data provides an additional check of statistical consistency. The energy of a configuration is proportional to the logarithm of its probability (in addition, because the entropy *S* is obtained from the cluster expansion, we can also compute the constant of proportionality). Concordance between the inferred and empirical energy distributions thus indicates that the real data could plausibly be generated from the inferred model. Figure 4 compares the data and model distributions of energies, showing that in most cases they closely overlap. A small discrepancy is introduced in PF00014 because of the reweighting procedure (here the histogram of the data is normalized by the sequence weights). The energy distribution for the lattice protein model is broader than for the data, though the peak is fit correctly. In contrast with models inferred using ACE, the distribution of energies of the data is less well reproduced with plmDCA (Supplementary Materials). The ability to estimate the probability of a configuration can be useful when comparing the likelihood of a configuration in two different models, for example to decide which family a given protein belongs to.

### 3.5 ACE accurately infers structural contacts for PF00014

In Fig. 6 we use the inferred couplings to predict pairs of residues that are in contact in the folded protein structure for PF00014, and we compare results from ACE to the standard contact prediction methods DCA [6] and plmDCA [19]. In this case the pairs of sites for which the Frobenius norm of the couplings is largest, including the average product correction (APC, see [33]), are predicted to be most likely to be in contact. We define contact residues to be those that are within 6Å of each other in the folded structure of the protein, and we exclude trivial contact pairs along the protein backbone (*i* – *j* ≤ 4).

**Figure 6:**
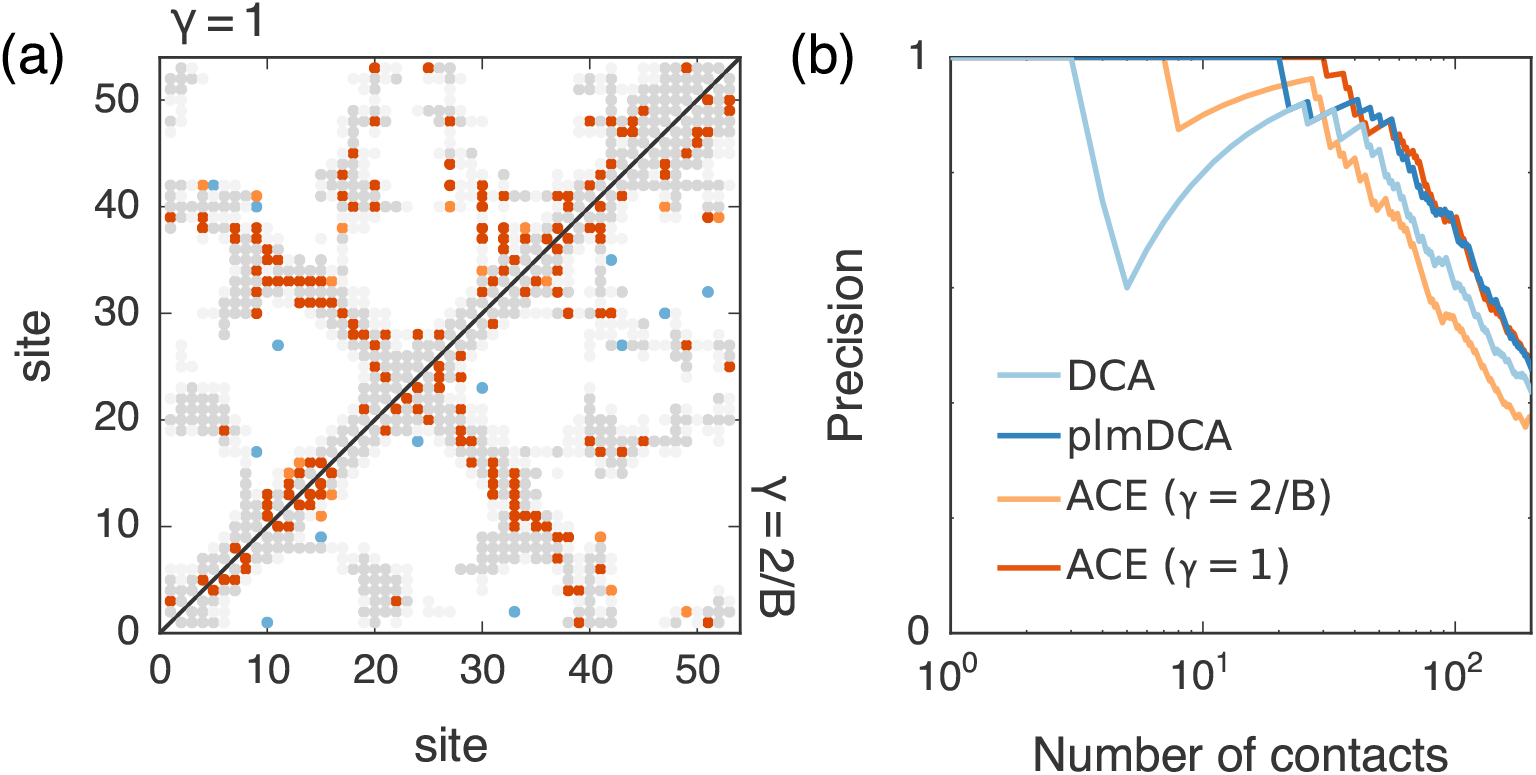
(a) Contact map for PF00014 inferred by ACE. Here we show the top 100 predicted contacts, with true predictions in orange and false predictions in blue. Other contact residues in the crystal structure are shown in gray. For true positives and other contact residues, close contacts (< 6Å) are darkly shaded and further contacts (< 8Å) are lightly shaded. The upper and lower triangulär parts of the contact map give predictions for the inferred model with strong regularization/no compression (γ = 1) and weak regularization/high compression (γ = 2/B), respectively. (b) Precision (ratio between the number of true predictions and the total number of predictions) as a function of the number of predictions for close contact residues that are widely separated on the protein backbone (i – j > 4). Results using ACE compare favorably with those from DCA [6] and are competitive with those from plmDCA [19].

The accuracy of contact predictions with ACE can be increased by decreasing the compression (*p*_o_ = 0) and using a large regularization (γ = 1), in the same spirit as the strong regularization employed in typical DCA and plmDCA approaches. Here we gauged the parameters for the least frequently observed amino acids to zero and computed the Frobenius norm of the couplings in the zero sum gauge (as is typical in DCA). The couplings are then strongly damped by regularization and the cluster expansion converges for maximal cluster sizes much smaller than those needed in the case with weaker regularization. Figure 6b shows that the precision in this case is competitive with the one obtained from plmDCA, and the prediction of the first ~ 30 contacts is slightly better for ACE. However, in this case we note that because of the small values of the couplings the generative properties of the inferred model are lost (see Supplementary Materials for the statistical fit of the model).

## 4 Discussion

Potts models have been successfully applied to study a variety of biological systems. However, the computational difficulty of the inverse Potts problem, i.e. the inference of a Potts model from correlation data, has presented a barrier to their use. Here we presented ACE, a flexible, easy-to-use method for solving the inverse Potts problem, which can be applied to analyze a wide variety of real and synthetic data. We also provide tools for automatically generating correlation data from multiple sequence alignments (MSA), making the analysis of this type of data even more accessible.

Here we have adapted the complexity of the inferred Potts models to the level of the sampling in the data. This is achieved by regrouping less frequently observed Potts states into a unique state (according to a threshold on entropy or frequency), then by a sparse inference procedure that omits interactions that are unnecessary for reproducing the statistics of the data to within the error bounds due to finite sampling. On artificial data we verified that compression of the number of Potts states allows a faster and more precise inference of the uncompressed model parameters while reducing overfitting. The methods of compression that we describe here can also be applied to other inference methods (including, for example, the DCA and plmDCA approaches discussed above), a topic of future study. In addition, as described above ACE yields sparser models when sampling is poor, leading to more robust inference.

This method allows for the simple construction of models from various types of data, and which can then be used to predict the evolution of experimental systems and their response to perturbations. Previous work has demonstrated promising applications of such models in a variety of different biological contexts. In neuroscience, the analysis of multi-electrode recordings has led to models that identify cell assemblies, which are thought of as basic units of memory [26]. Studies of MSAs of protein families allows for the prediction of pairs of residues in contact in the folded protein structure, giving insights on the protein structure from sequence information alone. Classical protein folding algorithms can be then used to refine the structure from contact predictions [15, 16, 17]. Potts models have also been used to describe the mutational landscape of viral and bacterial proteins, where they provide information about the effects of mutations on protein function, which could potentially be exploited to improve vaccine design and drug treatment [7, 8, 27, 9]. Recent work has also shown that a Boltzmann machine learning algorithm can be constructed to give a good generative model predicting the structure and functional dynamics of proteins [23]. Running such algorithms from a good initial guess of parameters, such as those obtained by ACE, could help to accelerate the inference procedure.

In the present work we have compared ACE with standard maximum entropy inference methods based on Gaussian and pseudo-likelihood approximations. These methods are particularly fast and adapted to find structural contacts and use, respectively, large pseudocounts and regularizations. Inference with ACE is generally slower than mean-field and pseudo-likelihood approaches. However, it allows for the accurate inference of underlying model parameters (when they are known), and for the construction of good generative models of the data when using a Bayesian value of the regularization strength (γ ≈ 1/*B*). In analogy with DCA and plmDCA, when using ACE with little compression (e.g. *p*_o_ = 0) and strong regularization the contact prediction obtained using traditional contact estimators is improved while the generative power of the inferred model is degraded.

An additional advantage of ACE is that it evaluates the entropy of the Potts model corresponding to a given set of data. For protein sequence data, this entropy gives a measure of the variability of the sequences in the same protein family, and can be used to predict site-dependent variability and robustness with respect to mutations [34]. We have now successfully applied the method to protein sequences of a few hundred amino acids in length collected from phylogenetically distant organisms, or longer sequences (up to 500 amino acids) for more phylogenetically related and less variables HIV MSA alignments.

## Acknowledgements

This work originates from the development of ACE in the Ising case in collaboration with R. Monasson, to whom we are grateful for many helpful discussions. We also thank D. Murakowski for his contribution to the development of the partition function expansion, and U. Ferrari and H. Jacquin for useful discussions.

